# Cross-reactive single-domain antibodies to hemagglutinin stem region protect mice from group 1 influenza A virus infection

**DOI:** 10.1101/2022.09.29.510074

**Authors:** Darya V. Voronina, Alina S. Bandelyuk, Alina Sh. Dzharullaeva, Olga Popova, Vladislav Yu. Kan, Ilias B. Esmagambetov, Irina A. Favorskaya, Dmitry V. Shcheblyakov, Boris S. Naroditskiy, Aleksandr L. Gintsburg

## Abstract

The continued evolution of influenza viruses reduces the effectiveness of vaccination and antiviral drugs. The identification of novel and universal agents for influenza prophylaxis and treatment is an urgent need. We have previously described two potent single-domain antibodies (VHH), G2.3 and H1.2, which efficiently neutralize H1N1 and H5N2 influenza viruses *in vivo*. In this study, we modified these VHHs with Fc-fragment to enhance their antiviral activity. Reformatting of G2.3 into bivalent Fc-fusion molecule increased its *in vitro* neutralizing activity against H1N1 and H2N3 viruses up to 20-fold and, moreover, resulted in obtaining the ability to neutralize H5N2 and H9N2 subtypes. We demonstrated that a dose as low as 0.6 mg/kg of G2.3-Fc or H1.2-Fc administered systemically or locally before infection could protect mice from lethal challenges with both H1N1 and H5N2 viruses. Furthermore, G2.3-Fc reduced the lung viral load to an undetectable level. Both VHH-Fc showed *in vivo* therapeutic efficacy when delivered via systemic or local route. The findings support G2.3-Fc as a potential therapeutic agent for both prophylaxis and therapy of Group 1 influenza A infection.

## Introduction

Influenza remains one of the major burdens in global healthcare, leading to 3 – 5 million severe illnesses and up to 650 000 lethal cases annually [1]. Annual seasonal epidemics are commonly caused by influenza virus types A and B, with a predominance of influenza A viruses (IAV). The high prevalence among the human population and the ability to generate pandemics that take millions of human lives make IAV of high epidemiological importance. The repeated seasonal epidemics and the chance of unpredictable pandemic outbreaks are associated with the antigenic evolution of IAV, which facilitates the escape from pre-existing immunity induced by prior infection or vaccination, and results in antiviral drug resistance and reduces the effectiveness of vaccination [2–8]. Thus, there is an urgent need in developing novel and effective antiviral agents for the treatment and prophylaxis of IAV infection.

Hemagglutinin (HA) is a surface glycoprotein, which ensures attachment and viral entry into the host cell, and is one of the primary targets for immune response. HA consists of two structurally and functionally distinct domains, variable globular head and conserved stem domain. The immunodominant globular head region is very susceptible to accumulating mutations under the pressure of an antibody response [9–14], and many of these antibodies are strain-specific. However, antibodies directed against the HA stem can bind a broad range of influenza subtypes, but are not usually produced in high titers [15–17]. Therefore, the HA stem domain is an attractive target for both specific prophylaxis and therapy with the potency to obtain an agent with broad-spectrum specificity.

Monoclonal antibodies (mAbs) have been proven as effective therapeutics for both non-infectious and infectious diseases [18–21]. Among different types of antibodies, camelid- or shark-derived single-domain antibodies (sdAbs) have several advantages for treating respiratory infections, including influenza virus infection, compared with conventional mAbs. sdAbs represent variable fragments of heavy-chain antibody, which recognize the antigen with only one domain, named VHH. Previous studies have reported the development of promising therapeutic VHH to various influenza antigens [22–26], in particular broadly reactive anti-HA stem VHHs [27–30]. Due to their unique CDR3 region, forming an extended loop, VHH have superiority over conventional immunoglobulins in binding hidden epitopes that may not be accessible for mAbs [31,32]. Nevertheless, their small size results in a short half-life *in vivo*, which may be a disadvantage in long-term therapeutic or prophylactic applications [33]. Moreover, an important role in limiting the development of influenza infection is played by the effector functions of antibodies mediated by the Fc-fragment, antibody-dependent cellular cytotoxicity (ADCC) and antibody-dependent cellular phagocytosis (ADCP) [34–40]. Hence, VHH is a promising tool for influenza therapy, but it is of great importance to optimize the structure of sdAbs to extend their half-life and obtain effector functions through modification with Fc-fragment.

Most influenza therapeutic mAbs are administered via systemic routes, typically intravenously, and in the case of animal studies, intraperitoneal (i.p.) administration is the predominant mode of delivery. However, the main targets for influenza virus and the main affected area are upper and lower respiratory tracts [41]. Studies have shown that administration of mAbs, including VHH and their modifications, via the local routes is effective and even improves the potency of mAbs in contrast to systemic delivery [42–46]. Therefore, direct delivery of mAbs to the airway may be an opportune approach for treatment and prevention of infection caused by influenza viruses.

In this study, we report two anti-HA stem sdAbs fused with Fc-fragment, G2.3-Fc and H1.2-Fc, which efficiently neutralize Group 1 influenza A viruses *in vitro* and *in vivo*. G2.3-Fc compared with H1.2-Fc exhibited a broader *in vitro* neutralizing activity and protected mice against H1N1 and H5N2 lethal challenge *in vivo* with greater potency. Our findings support G2.3-Fc as a potential agent for both prophylaxis and treatment of Group 1 IAV infection.

## Materials and methods

### Viruses and recombinant proteins

A/California/07/2009 (CA/09(H1N1)), A/Victoria/2570/2019 (VA/19(H1N1)), A/Duck/mallard/Moscow/4970/2018 (duck/MW/18(H1N1)), A/Mallard duck/Pennsylvania/10218/84 (duck/PA/84(H5N2)), A/Swine/Hong Kong/9/98 (swine/HK/98(H9N2)) and A/Black Duck/New Jersey/1580/78 (duck/NJ/78(H2N3)) were grown in embryonated chicken eggs according to standard viral culture techniques. The mouse adopted (ma) A/California/07/2009 (CA/09(H1N1)ma) and A/Mallard duck/Pennsylvania/10218/84 (duck/PA/84(H5N2)ma) were obtained by serial lung-to-lung passages as described elsewhere [47]. For ADCC and ADCP assays, recombinant serotype 5 adenovirus, Ad5-swH1opt/California, carrying a codon-optimized gene encoding HA A/California/07/2009 (H1N1) protein, was gently provided by Dr. Maxim M. Shmarov (N.F. Gamaleya NRCEM, Moscow, Russia). Recombinant full-length HA (rFL HA) subtype H1 (A/California/04/2009) was purchased from Sino Biological (Beijing, China).

### Cell lines

CHO-S cell line was obtained from Thermo Fisher Scientific (Waltham, MA, USA) cat. no. R80007. Caco-2 and A549 cell lines were purchased from the Russian collection of vertebrate cell lines (Saint Petersburg, Russia). Jurkat-Lucia™ NFAT-CD16 cells and Jurkat-Lucia™ NFAT-CD32 were obtained by InvivoGen (San Diego, CA, USA) cat. code jktl-nfat-cd16 and jktl-nfat-cd32, respectively.

### VHH-Fc fusion construction, expression and purification

Nucleotide sequences coding the previously described VHH, H1.2 and G2.3, [48] fused with human IgG1 Fc-fragment and Llama IgG2b hinge were purchased from Evrogen (Russia). CHO-S cells were transfected with VHH-Fc containing construct using the CHO Gro System (Mirus Bio, WI, USA), according to the manufacturer’s protocol. Expressed antibodies were purified from culture supernatant using HisTrap™ MabSelect™ Sure columns (Cytiva, Danaher Corporation, Washington, D.C., USA). Purity was analyzed by sodium dodecyl sulfate polyacrylamide gel electrophoresis (SDS–PAGE).

As positive controls for ELISA, MN, ADCC and ADCP reporter assays, we used broadly neutralizing VHH, SD38 and SD36, fused with hIgG1-Fc [27]. These VHH-Fc were synthesized, expressed, and purified as described above.

### ELISA

Polystyrene microplates (Corning Inc., NY, USA) were coated overnight at 4°C with 1 μg/ml rFL HA in carbonate-bicarbonate buffer pH 9.6, then wash three times with PBS containing 0.05% Tween-20 (TPBS) and incubated 1 h with blocking buffer (TPBS containing 1% w/v casein) at 37°C. After that, plates were rinsed three times with TPBS and VHH-Fc (serial dilutions in blocking buffer) were added to the wells and incubated for 1 h at 37°C. Immunoplates were washed four times and bound antibodies were detected using polyclonal goat anti-human IgG HRP-conjugated antibodies (MilliporeSigma, Burlington, MA, USA) diluted 1:20000 in blocking buffer. After five washes 3,3’5,5’-tetramethylbenzidine (TMB) (Bio-Rad, USA) was added as a substrate. Fifteen minutes later, the reaction was stopped by the addition of 1 M H2SO4 and the absorbance was read at 450 nm. The half maximal effective concentration (EC50) values were calculated using four-parameter logistic regression using GraphPad Prism 7 (GraphPad Software Inc., USA).

The antibody binding activity to untreated and denatured HA was measured as described above. rFL HA subtype H1 was denatured with 0.1% SDS, 50 mM DTT and a dry block thermostat at 99 °C for 10 min.

### *In vitro* HA cleavage assay

The procedure was performed as described previously [49], with minor modifications. rFL HA H1 (0.5 μg) was incubated with VHH-Fc (0.5 μg) for 1 h at room temperature and then 50 ng of N-tosyl-L-phenylalanine chloromethyl ketone treated trypsin (TPCK-trypsin) was added. The mixture was incubated for 10 min at 37°C. The samples were heated with Laemmli sample buffer with 2-mercaptoethanol at 95°C for 20 min and separated on SDS-PAGE using Any kD™ Mini-PROTEAN^®^ TGX™ Precast Protein Gels (Bio-Rad, USA).

### Low-pH-induced conformational change ELISA

The procedure was carried out as described previously, with minor modifications [49]. 1 μg/ml rFL HA in carbonate-bicarbonate buffer pH 9.6 was immobilized on the surface of 96-well polystyrene microplates (Corning Inc., NY, USA) overnight at 4°C. The plates were washed three times with TPBS and incubated for 1 h with blocking buffer (TPBS containing 1% w/v casein) at 37°C and 350 rpm. After that, plates were rinsed three times with TPBS. TPCK-treated trypsin (25 ng/ml) was added, after incubation for 1 h at 37°C and 350 rpm plates were washed again. Citrate buffer 0.1 M (with addition of 150 mM NaCl) with varying pH (7.4, 6 and 5) was added to the wells and immunoplates were incubated for 1 h at room temperature without shaking. Plates were washed thrice and VHH-Fc diluted in blocking buffer were added. The conventional ELISA protocol was performed as described above.

### Microneutralization (MN) assay

H1.2 and G2.3 in monovalent VHH format for MN assays were expressed and purified as described previously [48]. Caco-2 cells were maintained in complete Dulbecco’s Modified Eagle Medium (DMEM) supplemented with 20% fetal bovine serum (FBS). Caco-2 cells were seeded in 96-well culture plates (4*10^4^ cells/well) on the day before the experiment. Two-fold serially diluted VHH or VHH-Fc were mixed with an equal volume of virus (100 TCID50/well) in Medium 199 with Hanks’ salts (MilliporeSigma, USA) and incubated at 37 °C for 40 min. Caco-2 cells were washed twice with DMEM without FBS, samples were added and incubated with the cells at 37°C for 40 min. After incubation, the medium was replaced with a fresh medium and plates were incubated in a 5% CO2 incubator at 37°C for 96 h. The assay was performed in quadruplicate. Each 96-well plate contained positive (virus-inoculated cells) and negative (mock-inoculated cells) controls, as well as the control SD38-Fc. Neutralization of the virus was confirmed with hemagglutination inhibition assay. The cell medium was mixed with an equal volume of 0.75% Chicken erythrocytes and incubated for 30 min at room temperature. Negative (PBS with erythrocytes) and positive (8 hemagglutinating units of virus with erythrocytes) controls were included in every plate. Absence in button formation was scored as inhibition of hemagglutination. The minimal neutralizing concentration was defined as the lowest antibody concentration that completely inhibited agglutination of erythrocytes. Half-maximal inhibitory concentration (IC50) values were calculated using four-parameter logistic regression using Prism 7 (GraphPad Software Inc., USA).

### In vitro ADCC and ADCP Reporter Assay

A549 cells were maintained in DMEM supplemented with 10% inactivated FBS. A549 cells were seeded in 96-well culture plates 2*10^4^ cells/well on the day of the experiment. After the cell adhesion, culture media was replaced with the medium containing Ad5-swH1opt/California (15 PFU/cell) and plates were incubated in a 5% CO2 incubator at 37°C for 72 h. After incubation, the cells were washed once with a fresh medium and VHH-Fc diluted in the same medium were added to the wells. The plates were incubated in a 5% CO2 incubator at 37°C for 1 h, then the cells were washed once to remove unbinding VHH-Fc. The reporter cells Jurkat-Lucia™ NFAT-CD16 cells or Jurkat-Lucia™ NFAT-CD32 were added and the experiment was performed as described previously [50]. The analysis included positive control (SD38-Fc), negative control (non-relevant VHH-Fc) and intact cells. The assay was performed in quadruplicate.

### Prophylactic and therapeutic efficacy studies in mice

All animal procedures were approved by the Institutional Ethics Committee on Experimental Animals (protocol #19 dated 02.03.2022). Specific-pathogen–free female 6-8-week-old BALB/c mice were inoculated intranasal with 5 LD50 of mouse-adapted A/California/07/2009 (H1N1) or A/Mallard duck/Pennsylvania/10218/84 (H5N2) (50 μl, divided equally between the nostrils) and observed daily for clinical signs and weighed. Antibodies were diluted in phosphate-buffered saline (PBS) and administered intranasal or intraperitoneal in a volume of 50 μl or 200 μl respectively. Mice were anesthetized by inhaled anesthetic isoflurane before intranasal administration of VHH-Fc or challenge virus. Depending on the experiment, antibodies were administered up to 24 h before or after challenge, as specified in the figure legends. A 25% loss in body weight was determined as the clinical endpoint at which agonizing mice were humanely euthanized.

To determine the effect of antibody delivery on lung virus titer production, mice were administered i.p. with 3 mg/kg VHH-Fc, then, 24 h later challenged with 5 LD50 of virus and euthanized 4 days after challenge. Lung homogenates were prepared in 1 ml Medium 199 with Hanks’ salts, cleared by centrifugation at 4°C, and used for virus titration, as described above. Lung viral titers were expressed in TCID50 per milliliter and determined using the Reed and Muench method [51].

### Multiple sequence alignment and phylogenetic tree

Sequences of the FL HA protein of CA/09(H1N1), VA/19(H1N1), duck/MW/18(H1N1), duck/PA/84(H5N2), duck/NJ/78(H2N3) and swine/HK/98(H9N2) were downloaded from the Influenza Research Database (IRD) or the Nucleotide database (NCBI) and aligned using the MEGA X Software (Penn State University, USA) with the MUSCLE method. The phylogenetic tree was produced using the maximum likelihood method and visualized in the MEGA X program.

### Statistical analysis

All statistical analyses and plot generation were conducted using Prism 7 Software (GraphPad Software, Inc., USA). EC50 and IC50 values were calculated using four-parameter logistic regression. Differences between mice groups were analyzed using the nonparametric Kruskal-Wallis test (one way ANOVA), followed by Dunnett’s multiple comparisons test. The Mantel-Cox log-rank test was used to assess the statistical significance of Kaplan-Meier survival curve comparisons.

## Results

### Construction and characterization of VHH-Fc

In a previous study, we described anti-HA stem single-domain antibodies, G2.3 and H1.2, which showed potent neutralizing activity *in vivo* against H1N1 and H5N2 IAV [48]. In this study, we modified these VHH with human IgG1 Fc-fragment to obtain prolonged serum half-life required for prophylactic and therapeutic applications and to engage Fc-effector functions (Fig 1A). Purified antibodies were analyzed in SDS-PAGE, and the formation of dimeric Fc-fusion VHHs of approximately 85-90 kDa was confirmed (Fig 1B). The specific activity of VHH-Fc was determined in ELISA using rFL HA (A/California/04/2009) as antigen (S1 Fig).

**Fig 1.**
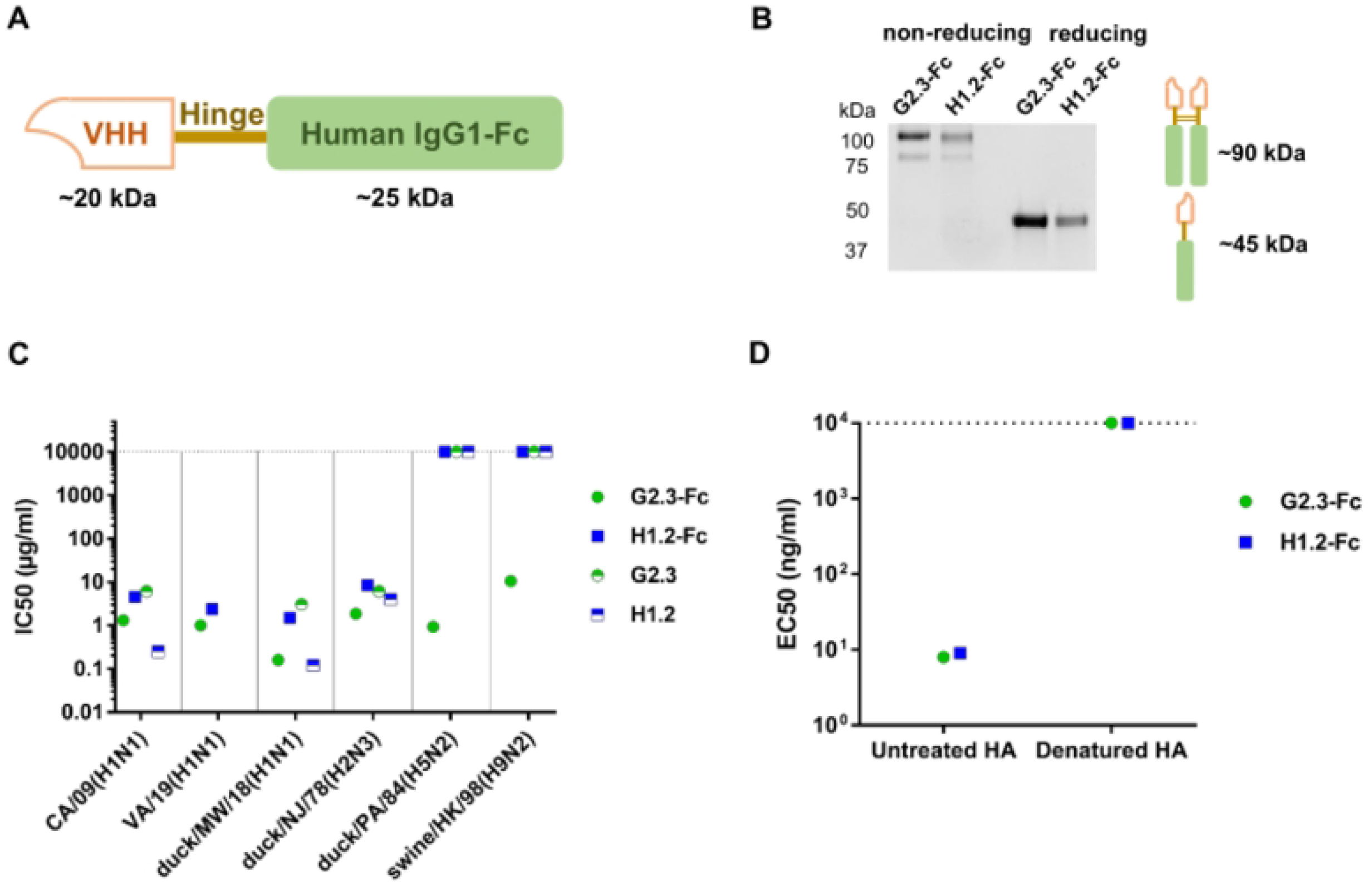
Characterization of G2.3-Fc and H1.2-Fc. **(A)** A schematic representation of the VHH-Fc constructs. **(B)** Analysis of purified VHH-Fc using SDS-PAGE. Theoretical molecular weights for VHH-Fc under reducing and non-reducing conditions are ~45 kDa and 85-90 kDa, respectively. **(C)** Neutralizing activity of VHH and VHH-Fc *in vitro* against Group 1 IAV. IC50 is expressed in microgram per milliliter and was measured in quadruplicate. **(D)** EC50 values determined in ELISA showing binding of VHH-Fc to untreated and denatured rFL HA (A/California/04/2009). EC50 values are expressed in nanogram per milliliter. Dotted lines indicate the absence of activity.

To determine the neutralizing potency of our antibodies, we performed a MN assay with H5N2, H2N3, H9N2 influenza viruses, and different strains of H1N1 subtype (Fig 1C, S2 Fig). We compared the activity of Fc-fused and monovalent forms of VHH to explore whether dimerization of VHH affected their neutralizing performance. We observed that G2.3-Fc, compared to its monovalent form, not only improved potency for H1 and H2 subtypes but also obtained the ability to neutralize H5 and H9 subtypes. It was shown that in contrast to G2.3-Fc antibody H1.2-Fc exhibited lower neutralizing potency against H1 and H2 viruses and did not neutralize H5 and H9 viruses *in vitro*. Interestingly, reformatting H1.2 VHH into a bivalent format decreased its *in vitro* potency. We speculate that it might be a result of steric hindrance [52,53] and spatial interference probably results in a reduced neutralizing activity of VHH-Fc, compared to the small-sized VHH domain.

To define whether the binding of G2.3-Fc and H1.2-Fc to H1 HA protein was conformation-dependent we performed ELISA with denatured HA (Fig 1D). The antibodies lost their ability to bind to HA after denaturation, but maintained binding activity with native HA protein. ELISA studies showed that both VHH-Fc recognize conformational epitopes on the surface of HA protein.

To understand the potential mechanism underlying the neutralizing activity, we explored the influence of the antibodies on the membrane fusion process. This process consists of two stages, which can be affected by antibodies: HA cleavage by host-cell trypsin-like proteases and low-pH-induced conformational change of HA in endosomes. Initially, we modeled the proteolytic activation of HA with TPCK-trypsin in the presence of VHH-Fc. The results showed that both VHH-Fc, the same as control antibody SD38-Fc, could not prevent protease cleavage of HA (Fig 2A). Afterwards, to test whether G2.3-Fc and H1.2-Fc could bind to HA, which undergoes low-pH-induced conformational change, we subjected rFL HA H1 to proteolytic activation, then treated cleaved HA with low pH and performed conventional ELISA protocol. ELISA studies showed that both G2.3-Fc and H1.2-Fc maintain their binding activity at pH 7.4 and pH 6, while with decreasing pH to 5, the activity of VHH-Fc has slightly declined (Fig 2B and Fig 2C). The results obtained for G2.3-Fc and H1.2-Fc were comparable to those for the control VHH-Fc, SD38-Fc (S3 FigA). In contrast, the attachment of the control antibody SD36-Fc markedly decreased with a decrease in the pH (S3 FigB), which is consistent with the results obtained by the Laursen et al. [27]. We speculate that the mechanism of antiviral action of G2.3-Fc and H1.2-Fc may lies in inhibiting low-pH-induced conformational changes that would block membrane fusion.

**Fig 2.**
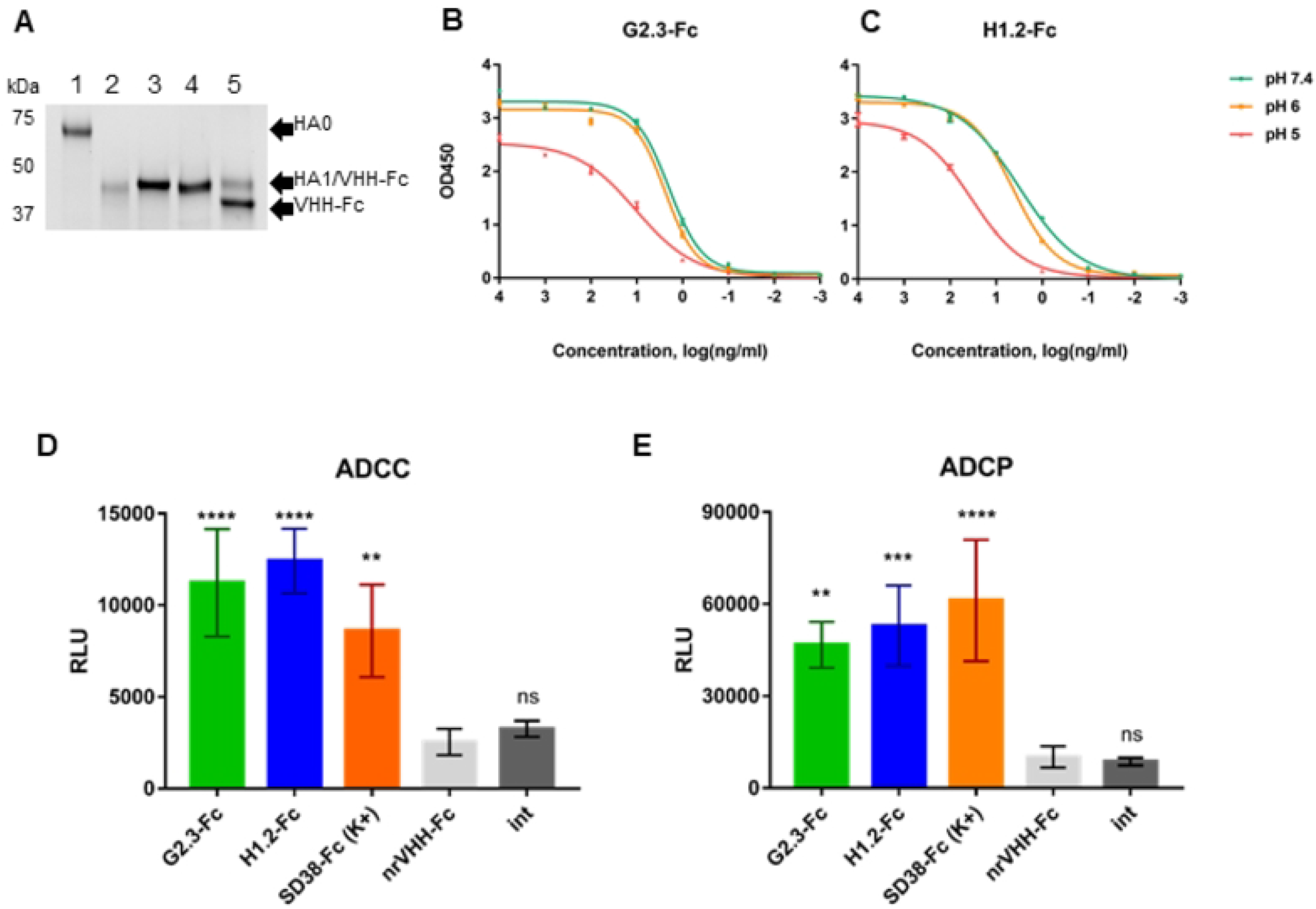
Mechanism of antiviral action of VHH-Fc. **(A)** Cleavage inhibition assay: rFL HA H1 was pre-incubated with VHH-Fc and then cleaved with TPCK-trypsin. 1 – rFL HA H1 (~65-67 kDa) under reducing conditions; 2 – product of rFL HA H1 tryptic digestion: HA1 (~45 kDa), HA2 band is not visible; 3 – rFL HA H1 preincubated with G2.3-Fc, HA1 band overlaps with the single chain of VHH-Fc band due to similar MW; 4 – rFL HA H1 preincubated with H1.2-Fc, HA1 band overlaps with the single chain of VHH-Fc band; 5 – rFL HA H1 preincubated with SD38-Fc, HA1 band separates from the single chain of VHH-Fc band. **(B) and (C)** ELISA with rFL HA H1 subjected to trypsinization and treated with different pH (citrate buffer pH 7.4, 6 and 5). **(D) and (E)** Evaluation of Fc-mediated functions, ADCC and ADCP: the effector cells, featuring CD16 (ADCC, **D**) or CD32 (ADCP, **E**) were added to A549 cells, expressing HA A/California/07/2009 (H1N1) protein and previously incubated with VHH-Fc. ADCC and ADCP activities were expressed as RLU – relative luminescence units. Error bars represent means ± SD. Asterisks indicate significant differences from the group, treated with non-relevant VHH-Fc (nrVHH-Fc) (****, P < 0.0001; ns, non-significant). For **D**: **, P = 0.0018; for **E**: **, P = 0.0012; ***, P = 0.0003. int – intact cells.

In the next stage, we evaluated the effector functions of VHH-Fc since previous studies indicate a protective role for Fc-mediated functions of antibodies against influenza infection *in vivo* [35–37,54]. To assess the possibility of FcγR activation by ours VHH-Fc, we performed ADCC and ADCP reporter assays. A549 cells were transduced with Ad5, carrying a gene encoding HA A/California/07/2009 (H1N1) protein. VHH-Fc were incubated with target cells for 1 h to bind to membrane-anchored HA. Subsequently, Jurkat-Lucia™ NFAT-CD16 cells or Jurkat-Lucia™ NFAT-CD32 effector cells (InvivoGen, USA) were added for further incubation and bioluminescence was measured. In both ADCC and ADCP tests activity of G2.3-Fc and H1.2-Fc was statistically significant compared to the non-relevant VHH-Fc and intact cells, and they showed similar to SD38-Fc potency (Fig 2D and Fig 2E). These data showed that G2.3-Fc and H1.2-Fc could effectively engaged Fc-effector functions.

### Prophylactic efficacy of VHH-Fc administered systemically or locally against H1N1 and H5N2 IAV

We initially evaluated the effect of systemic delivery for both antibodies. Mice were injected i.p. with 3 or 0.6 mg/kg of VHH-Fc 24 h before infection with 5 LD50 of CA/09(H1N1)ma or duck/PA/84(H5N2)ma virus. As shown in Fig 3, all the mice administered 3 mg/kg of G2.3-Fc survived both H1 and H5 virus challenge and did not demonstrate any weight loss. Mice that received H1.2-Fc (3 mg/kg) also survived H1 and H5 virus challenge, but exhibited transient weight loss with recovery by day 7 postinfection. In contrast, all control mice treated with the vehicle (PBS) lost weight and succumbed to infection or reached a humane endpoint (25% weight loss) and were euthanized by day 7 postinfection. Surprisingly, the administration of as low as 0.6 mg/kg of both VHH-Fc showed full protection against both viruses, antibody-treated mice exhibited no or transient weight loss (in case of H1.2-Fc prophylaxis) with recovery by day 9 postinfection. In the control groups, 4 of 5 and 5 of 5 mice, challenged with H1N1 and H5N2, respectively, succumbed to infection or reached the endpoint by day 8 postinfection.

**Fig 3.**
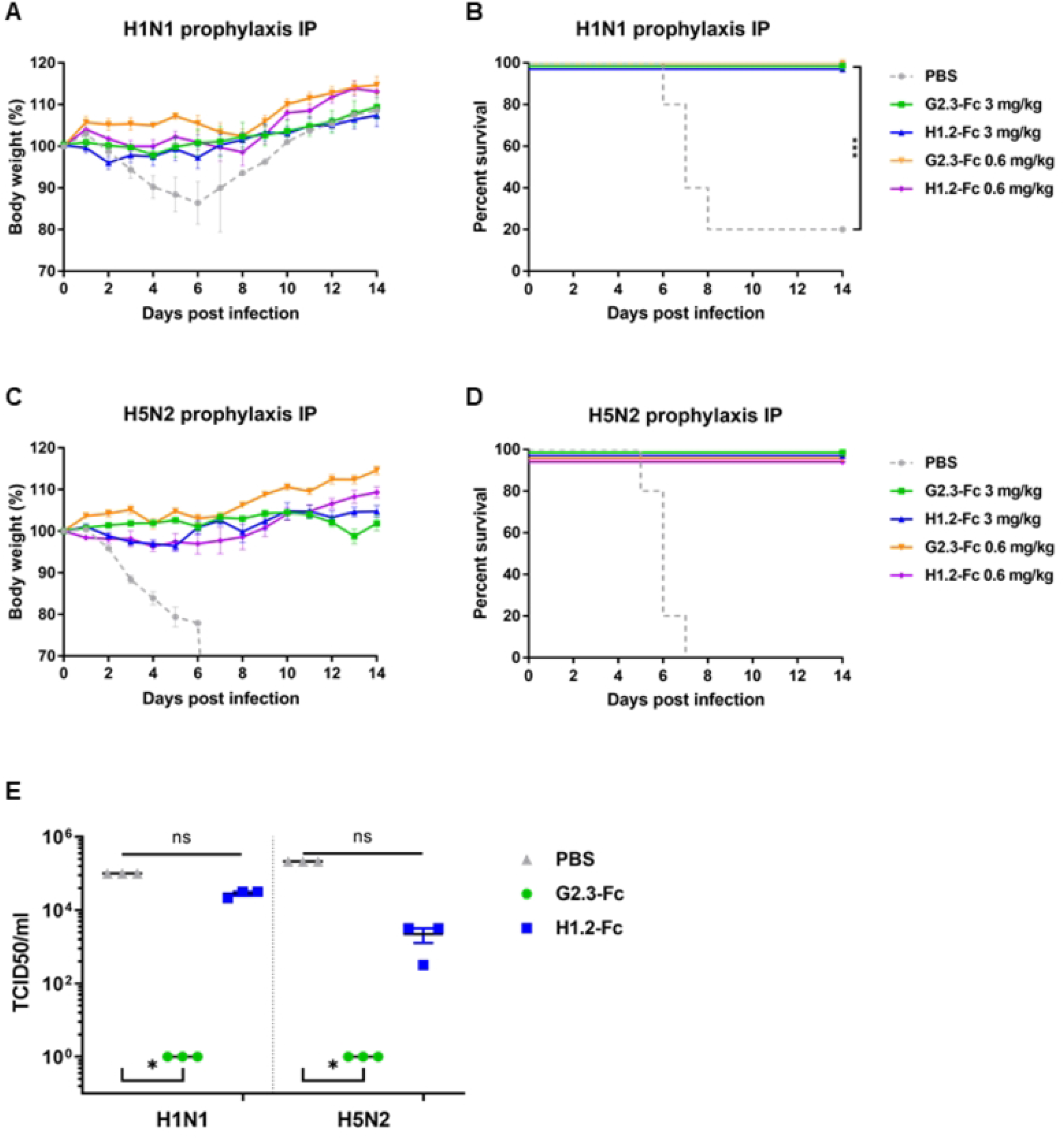
Prophylactic efficacy of systemically administered VHH-Fc in mice. Body weight changes and survival curves of BALB/c mice (n = 5 per group), treated via the i.p. route with G2.3-Fc or H1.2-Fc in different doses 24 h before infection with influenza CA/09(H1N1)ma virus **(A and B)** or duck/PA/84(H5N2)ma virus **(C and D)**. As a negative control, a vehicle (PBS) was administered 24 h before infection via the i.p. route. Body weight curves represent mean value ± SEM. Significant differences between groups are shown: ***, P = 0.0002. **(E)** Lung viral titers following G2.3-Fc or H1.2-Fc prophylaxis and H1N1 or H5N2 virus challenge in BALB/c mice. Lung viral titers are expressed in TCID50 per milliliter. Error bars represent means ± SEM. Asterisks indicate significant differences from the control group (*, P = 0.0105; ns – non-significant).

To evaluate the effect of systemic antibody delivery on lung virus titer production, mice were treated with 3 mg/kg of VHH-Fc 24 h before infection with 5 LD50 of CA/09(H1N1)ma or duck/PA/84(H5N2)ma virus. On day 4 postinfection mice were euthanized and lungs were harvested to determine viral titers. Prophylactic i.p. administration of G2.3-Fc reduced lung viral titers compared with the groups that received the vehicle (Fig 3E). The lung viral titers in all groups treated with G2.3-Fc were undetectable, whereas the administration of H1.2-Fc did not lead to a statistically significant reduction in H1N1 and H5N2 virus titers. Overall, the results are consistent with the absence of clinical symptoms in G2.3-Fc – treated mice.

Based on early studies that showed improved *in vivo* efficacy of airway antibody delivery compared with systemic delivery, we performed a prophylactic administration of VHH-Fc via the intranasal route [43,44]. Mice were initially treated 0.6 mg/kg of G2.3-Fc or H1.2-Fc and were then challenged 24 h or 1 h later with 5 LD50 of CA/09(H1N1)ma or duck/PA/84(H5N2)ma virus. The survival and weight loss results for mice challenged with H1N1 are shown in Fig 4A and Fig 4B. Mice treated with the vehicle control lost weight rapidly and 4 of 5 mice succumbed to infection or reached the endpoint by day 8. All mice treated with antibodies 1 h before H1N1 infection survived the lethal challenge without transient weight loss. Administration of VHH-Fc 24 h before infection resulted in reduced protection compared with those treated 1 h before challenge. Mice that were inoculated i.n. with G2.3-Fc 24 h before H1N1 challenge showed 80% survival rate and a weight loss of 5% of their initial body weight, while only 40% of mice that received H1.2-Fc survived. In the context of the H5N2 challenge (Fig 4C and Fig 4D), all PBS control animals lost weight and succumbed to infection by day 9. Mice treated with VHH-Fc 1 h before challenge were fully protected from death, weight loss and clinical symptoms. Nevertheless, when infection occurred 24 h after antibody treatment 80% mice survived, the difference between groups who received VHH-Fc 1 h and 24 h before infection was statistically insignificant (P > 0.05).

**Fig 4.**
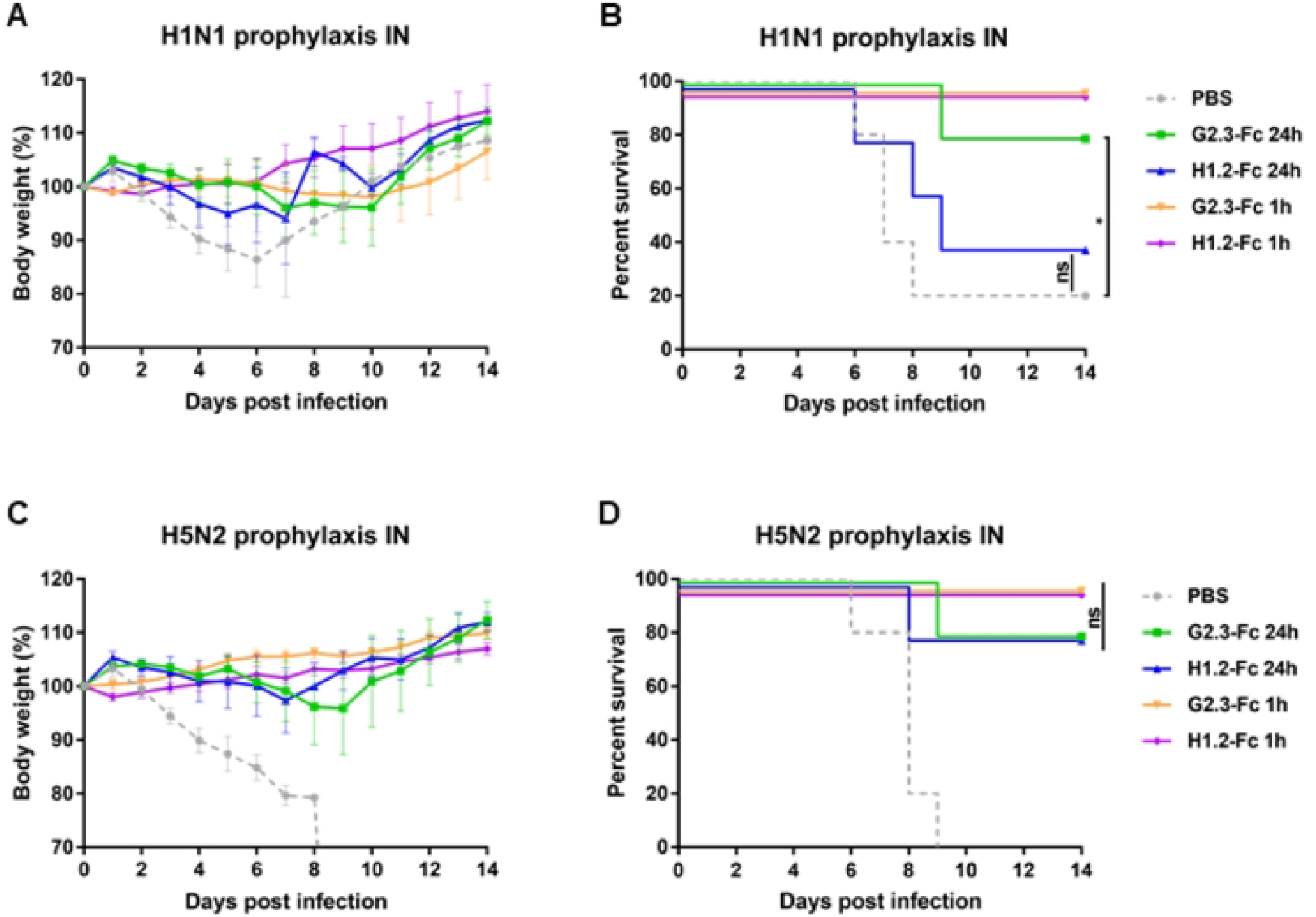
Prophylactic efficacy of locally administered VHH-Fc in mice. Body weight changes and survival curves of mice (n = 5 per group), treated via the i.n. route with G2.3-Fc or H1.2-Fc in dose of 0.6 mg/kg 24 h or 1 h before infection with influenza CA/09(H1N1)ma virus **(A and B)** or duck/PA/84(H5N2)ma virus **(C and D)**. As a negative control, a vehicle (PBS) was administered 1 h before infection via the i.n. route. Body weight curves represent mean value ± SEM. Significant differences between groups are shown: *, P = 0.033; ns – non-significant.

Thus, obtained results indicated that G2.3-Fc and H1.2-Fc could protect mice against H1N1 and H5N2 viruses *in vivo*. Prophylaxis with 0.6 mg/kg of G2.3-Fc is effective against both H1 and H5 viruses, while administered whether via the systemic or local route.

### Therapeutic efficacy of VHH-Fc administered systemically or locally against H1N1 and H5N2 IAV

After observing the efficacy of systemic and local delivery of VHH-Fc in a prophylactic regimen at low doses, we investigated the therapeutic effectiveness of G2.3-Fc and H1.2-Fc, when administered by different routes. Mice were challenged with 5 LD50 of CA/09(H1N1)ma virus or duck/PA/84(H5N2)ma virus and 2 h, 8 h, or 24 h later received 10 mg/kg of VHH-Fc via i.p. or 2 mg/kg via i.n. route. A control group was inoculated i.n. with the vehicle (PBS) 2 h postinfection.

While testing the therapeutic efficiency, G2.3-Fc could completely prevent symptom onset and death when administered 2 h (Fig 5A and 5B, Fig 6A, and 6B) or 8 h (Fig 5C and 5D, Fig 6C, and 6D) after the challenge with both H1 and H5 influenza viruses. We observed similar efficacy of G2.3-Fc at a dose of 2 mg/kg delivered locally and 10 mg/kg delivered systemically. Mice treated with G2.3-Fc had no signs of respiratory distress and exhibited no or transient weight reduction of only 3% of their initial body mass. In the context of H1.2-Fc, the results were similar to that obtained for G2.3-Fc, but a slightly greater weight loss was observed, up to 6%, when H1.2-Fc was administered 2 h postinfection with H5N2 virus (Fig 6A and B). Unfortunately, 24-h delay in a single administration of VHH-Fc after the challenge with H1N1 influenza virus did not show significant protection, although 2 of 5 mice had survived in the groups treated i.p. with 10 mg/kg G2.3-Fc and i.n. with 2 mg/kg H1.2-Fc (S4 FigA and B). All mice in the control and treated 24 h postinfection groups experienced rapid and strong weight loss, but survived mice regained or even exceeded their initial body weight.

**Fig 5.**
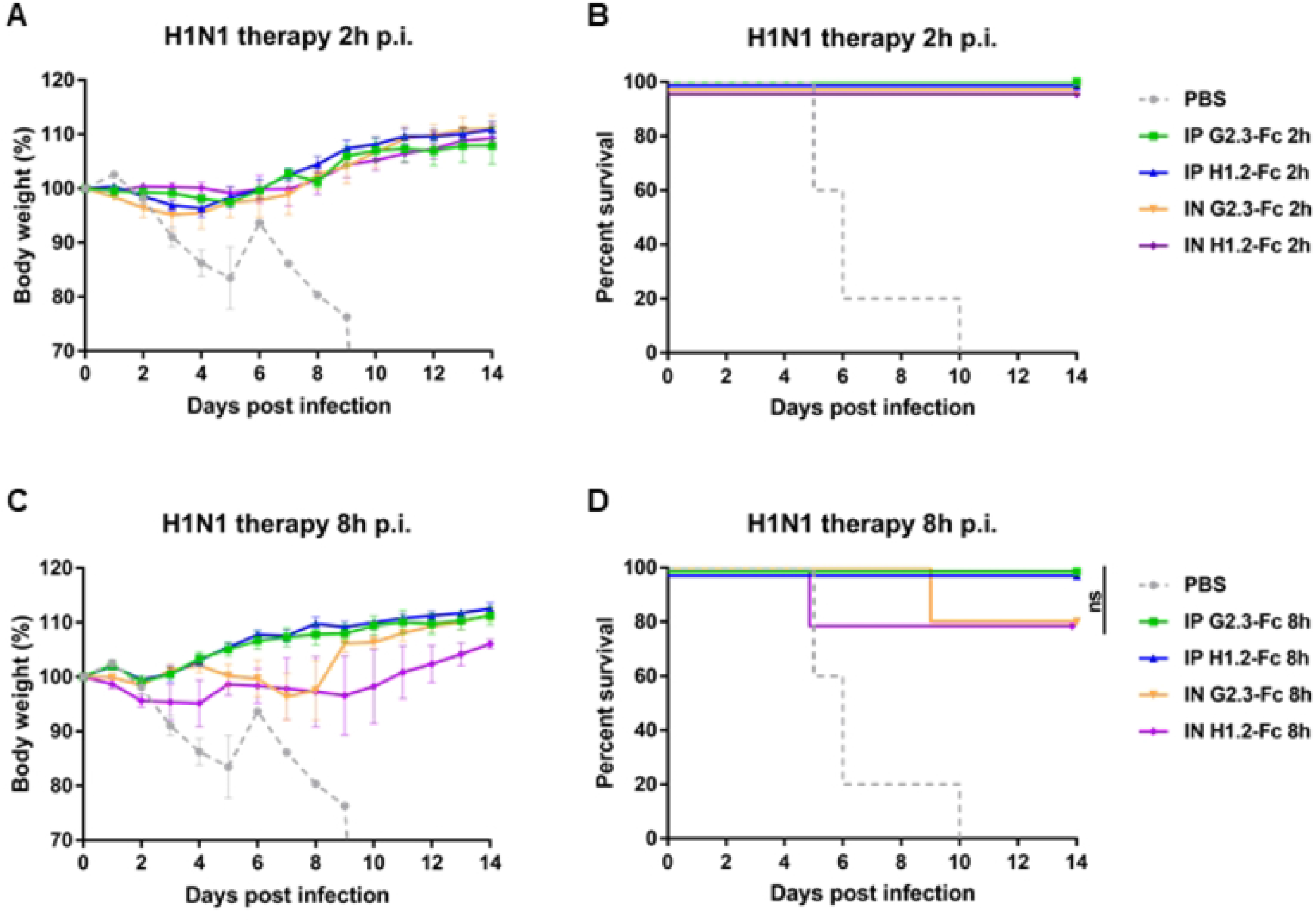
Therapeutic efficacy of VHH-Fc against H1N1 IAV in BALB/c mice. Body weights and survival curves of mice (n = 5 per group), treated with G2.3-Fc or H1.2-Fc with a dose of 10 mg/kg in case of i.p. injection or 2 mg/kg when delivered via the i.n. route. VHH-Fc were administered 2 h **(A and B)** or 8 h **(C and D)** after the infection with 5 LD_50_ of influenza CA/09(H1N1)ma virus. As a negative control, a vehicle (PBS) was administered 2 h after the infection via the i.n. route. Body weight curves represent mean value ± SEM. ns – the difference between groups is non-significant.

**Fig 6.**
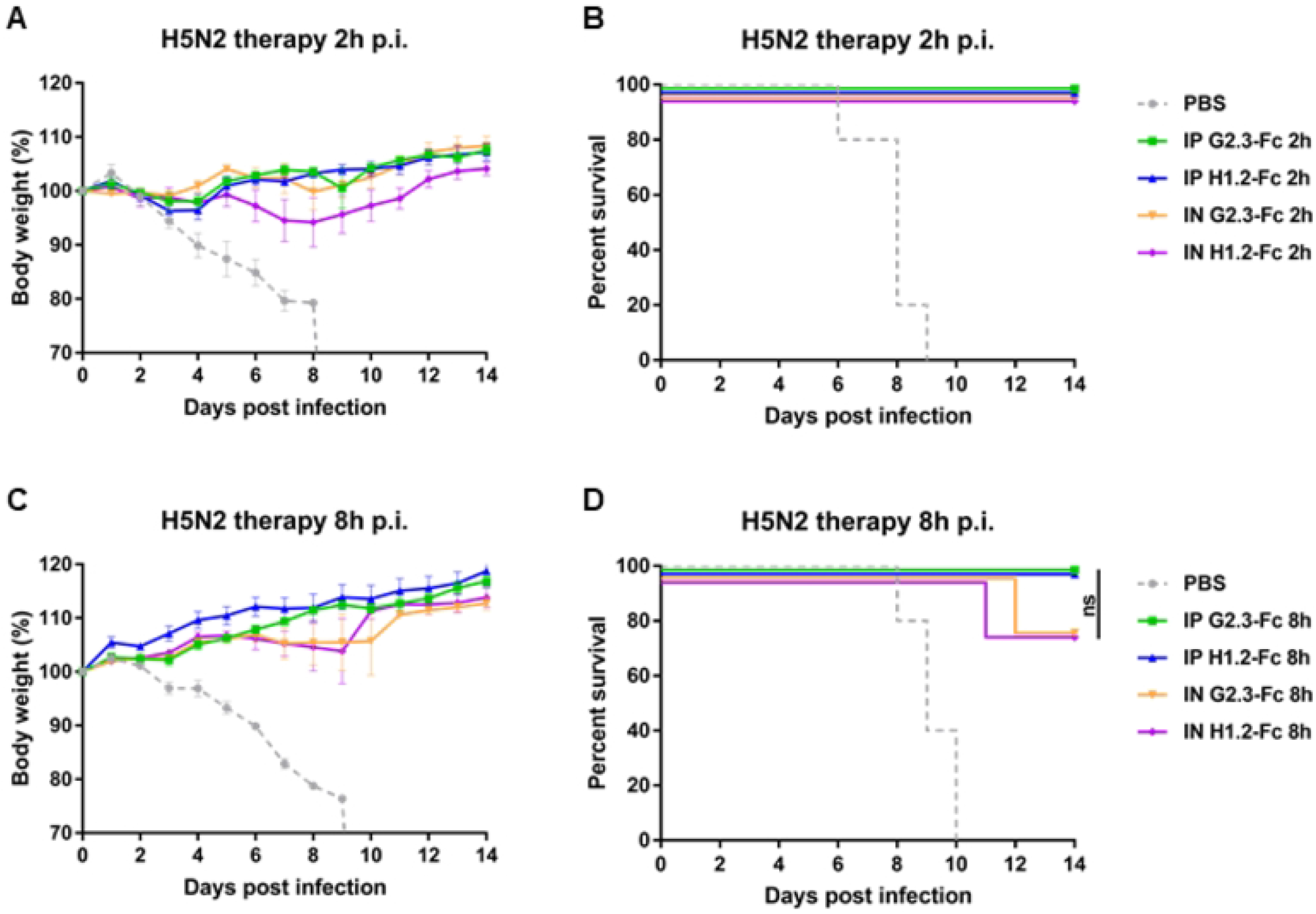
Therapeutic efficacy of VHH-Fc against H5N2 IAV in BALB/c mice. Body weights and survival curves of mice (n = 5 per group), treated with G2.3-Fc or H1.2-Fc with a dose of 10 mg/kg in case of i.p. injection or 2 mg/kg when delivered via the i.n. route. VHH-Fc were administered 2 h **(A and B)** or 8 h **(C and D)** after the infection with 5 LD50 of influenza duck/PA/84(H5N2)ma virus. As a negative control a vehicle (PBS) was administered 2 h (A and B) or 8 h (C and D) after the infection via the i.n. route. Body weight curves represent mean value ± SEM. ns – the difference between groups is non-significant.

Additionally, we tested multiple administration of VHH-Fc (6 mg/kg 24 h, 48 h and 72 h postinfection) via the i.p. route against both H1N1 and H5N2, and it improved G2.3-Fc antibody efficacy: 4 of 6 mice survived the challenge, although they could not completely regain initial body weight during the 2 weeks of observation (S4 FigC, D, E, and F).

Overall, these findings demonstrate that G2.3-Fc and H1.2-Fc could therapeutically protect mice from H1N1 and H5N2 IAV, furthermore i.n. administration of 2 mg/kg of VHH-Fc had the similar potency compared to the same VHH-Fc delivered via the i.p. route in a dose of 10 mg/kg.

## Discussion

influenza A viruses (IAV) lead to death of hundreds of thousands of people in annual epidemics worldwide and can initiate a pandemic, therefore remain a major public health problem. Among the Group 1 IAV, only H1 and H2 subtypes have caused pandemics and due to their rapid mutations in HA maintain this possibility [55,56]. Additionally, recent outbreaks of the H5 and H9 subtypes with high mortality rates made them a significant pandemic threat [57,58]. Thus, it is an urgent necessity to design the therapeutics against influenza infection and broadly neutralizing antibodies can be a useful tool for this purpose. Here, we describe a subtype cross-reactive VHH-Fc, which were generated based on our previously reported VHHs, G2.3 and H1.2, showed potent neutralizing activity *in vivo* against H1N1 and H5N2 IAV [48].

In the context of influenza viruses, antibody’s Fc-effector functions play a critical role in protection against infection and disease [36,37,59]. It has been reported that HA stem-specific cross-reactive antibodies provide Fc-dependent immunity, while strain-specific HA head antibodies protection is Fc-independent [35–37,39,54]. The inclusion of Fc-domain to the VHH structure not only adds Fc-effector functions, but also increases serum half-life [60,61] and improves binding ability of single-domain antibody by bivalency-imparting [28,62,63]. Fc-fragment of human IgG1 is the isotype of choice for mAb-based therapeutic, as it effectively binds to C1q and different Fc-receptors, thus provides a cell-mediated mechanisms, required for viral elimination. A recent study of broadly-neutralizing VHHs against influenza confirmed a significant increase in *in vivo* potency and neutralization breadth for Fc-fused VHHs [27]. In our work, two VHHs, H1.2 and G2.3, were linked to human IgG1-Fc to add them effector functions and retard their clearance.

The neutralization breadth of VHH-Fc was studied *in vitro* in microneutralization assay on Caco-2 cells. G2.3-Fc exhibited greater neutralizing activity compared to H1.2-Fc, being able to neutralize all tested subtypes of Group 1 IAV, including H1 strains isolated from different species, H2, H5 and H9. Fusion with Fc-fragment not only increased the neutralizing potency of G2.3-Fc against H1 and H2 subtypes, but also allowed the antibody to effectively inhibit growth of H5 and H9 subtypes compared to monovalent G2.3, which is consistent with previously published studies [27–29]. Surprisingly, converting of H1.2 into bivalent Fc-fusion form didn’t give the same results. We speculate that it might be a result of steric hindrance [52,53] and spatial interference probably results in a reduced neutralizing activity of VHH-Fc, compared to the small-sized VHH domain.

Many anti-stem broadly-neutralizing antibodies block viral entry by inhibition of the membrane fusion, thus providing the neutralizing effect *in vitro* [64–66]. Membrane fusion consists of two processes: proteolytic activation of HA and pH-induced conformational change in endosomes. Neither G2.3-Fc nor H1.2-Fc could prevent protease cleavage of HA. Though, given the fact that both G2.3-Fc and H1.2-Fc could bind to low-pH-treated conformation of H1 HA, we can assume that the possible antiviral mechanism of VHH-Fc lies in inhibiting low-pH-induced conformational changes. Identifying epitopes in the HA protein for our VHH-Fc could further shed the light on the mechanism of action.

The results of this study revealed that administration of both VHH-Fc can effectively protect mice against lethal H1N1 and H5N2 challenge, when administered either via a systemic or local route in prophylaxis and therapeutic regimens. We first assessed the prophylactic efficacy of VHH-Fc delivered via a systemic (i.p.) route. A prophylactic dose as low as 0.6 mg/kg of both G2.3-Fc and H1.2-Fc was sufficient to protect mice from disease and death with H1N1 or H5N2 IAV. Moreover, administration of G2.3-Fc reduced the viral loads in the lungs below limits of detection. Considering that the influenza viruses mainly affect the respiratory tract, it could be beneficial to deliver therapeutic antibodies directly to the site of infection [41]. The bioavailability of antibodies administered locally exceeds systemic administration, which allows reducing the amount of treatment and, accordingly, its cost [43,44]. The undeniable advantage of local administration of antibodies is not only a reduction in the overall dose, but additionally a more convenient application and diminishing the risks associated with intravenous drug delivery [67]. Therefore, our next step was to determine whether the prophylactic intranasal inoculation of VHH-Fc had the comparable protection against H1 and H5 IAV challenges as i.p. administration. We observed that 0.6 mg/kg of G2.3-Fc delivered 24 h or 1 h before infection was required for statistically significant protection of the mice challenged with either H1N1 or H5N2 IAV. Although we did not investigate the effect of the local administration of VHH-Fc in lower doses, according to previous studies, it is possible to improve the potency of locally administered antibody up to 10-fold greater that the same antibody exhibits, when delivered via systemic route [43,44].

Our study shows that both G2.3-Fc and H1.2-Fc can protect mice in the therapeutic mode when delivered via i.n. or i.p. route up to 8 h after the infection. The dose of 2 mg/kg delivered i.n. resulted in a similar protection as 10 mg/kg dose administered via systemic route. Although therapy with a single dose of VHH-Fc 24 h post-infection did not offer complete protection against H1N1 infection, survived mice gained weight and even exceed their initial body mass to the end of experiment. Three injections of G2.3-Fc on the first, second and third days after infection results in protection from dying for 60% of mice. An improvement in the therapeutic effectiveness of VHH-Fc can be achieved by increasing the amount of delivered antibody, due to multiple injections with a certain interval or by obtaining an antibody cocktail [68–72]. In future studies, we will explore whether these methods could enhance the potency of our VHH-Fc. Thus, we described two VHH-Fc, which effectively protect mice in prophylaxis and therapy from Group 1 IAV infection at low dosage when administered not only via systemic route, but also via local (i.n.) route. These results demonstrate great opportunities for prophylactic and therapeutic agents against influenza and others respiratory diseases.

Interestingly, H1.2-Fc could not ensure the neutralization of H5 subtype *in vitro*, notwithstanding the *in vivo* efficacy against H5 IAV. It should be noted, that protective potency of antibodies *in vivo* also depends on the Fc-mediated mechanisms including ADCC, ADCP and CDC [34,37,54,73]. Thus, antibodies lacking neutralizing activity *in vitro* can prevent infection *in vivo* by inhibiting virus release from infected cells and cell-to-cell spreading of viral progeny [74–78]. We have shown that both G2.3-Fc and H1.2-Fc have ADCC and ADCP activity *in vitro*, thus suggesting that G2.3-Fc and H1.2-Fc could engage Fc-effector functions for *in vivo* protection. Whether VHH-Fc could prevent cell-to-cell viral spread needs further investigation.

In conclusion, we developed two Fc-fused sdAbs (G2.3-Fc and H1.2-Fc) showed cross-protection against Group 1 IAV in prophylactic and therapeutic regimens. This study reported efficacy of local administration of antibodies on a par with systemic delivery in the context of influenza infection. The local passive immunotherapy via the mucosal route allows self-administration to patients, reduces the risks associated with injectable drug delivery. These VHH-Fc can find their clinical application as preventive treatment and as emergency therapy for IAV infection. The findings of this study could provide an effective strategy for influenza prevention and treatment.

## Author contributions

Writing – Original Draft Preparation: DV. Writing – Review & Editing: IF. Investigation: DV, AB, AS, VK, OP, IE. Formal Analysis: DV, AB, AS. Conceptualization: DV, IE, DS. Project Administration: DS. Supervision: DS, BN, AG.

---

**S1 Fig. Determining of specific activity of VHH-Fc in ELISA.**100 ng/well of rFL HA (A/California/04/2009) was used as antigen, EC50 was 8 ng/ml for G2.3-Fc and 9 ng/ml for H1.2-Fc.

**S2 Fig. (A) Summary of IAV used for MN and *in vivo* experiments.** ma – mouse adapted. **(B) IC50 values of VHH-Fc and their monomeric form.** MN analysis was performed on Caco2 cell line in quadruplicate. nt – not tested, NN – non-neutralize. **(C) Phylogenetic analysis of IAV used for MN in this study**. Sequences of the FL HA protein were and aligned using the MEGA X Software (Penn State University, USA) with MUSCLE method. The phylogenetic tree was produced using the maximum likelihood method and visualized in the MEGA X program.

**S3 Fig A and B. Low pH-induced conformation change ELISA.** ELISA with rFL HA H1 subjected to trypsinization and treated with low pH (citrate buffer pH 7.4, 6 and 5). **(A)** SD38-Fc; **(B)** SD36-Fc.

**S4 Fig A and B. Therapeutic efficacy of VHH-Fc against H1N1 IAV in BALB/c mice.** Body weights **(A)** and survival **(B)** curves of mice (n = 5 per group), treated with G2.3-Fc or H1.2-Fc 24 h postinfection with a dose of 10 mg/kg in case of i.p. injection or 2 mg/kg when delivered via the i.n. route. As a negative control, a vehicle (PBS) was administered 2 h after the infection via the i.n. route. Body weight curves represent mean value ± SEM. ns – the difference between groups is non-significant. **C, D, E, and F. Therapeutic efficacy of multiple administration of VHH-Fc.** Body weights **(C and E)** and survival **(D and F)** curves of mice (n = 6 per group), treated with G2.3-Fc or H1.2-Fc 24 h, 48 h, and 72 h postinfection with a dose of 6 mg/kg each time via the i.p. route. As a negative control, a vehicle (PBS) was administered 24 h, 48 h and 72 h after the infection via the i.p. route. Body weight curves represent mean value ± SEM. Significant differences between groups are shown: D: **, P = 0.0072; F: **, P = 0.0038; ns – non-significant.

## Notes

### Competing Interest Statement

The authors have declared no competing interest.

## References

1. Iuliano AD, Roguski KM, Chang HH, Muscatello DJ, Palekar R, Tempia S, Cohen C, Gran JM, Schanzer D, Cowling BJ, Wu P, Kyncl J, Ang LW, Park M, Redlberger-Fritz M, Yu H, Espenhain L, Krishnan A, Emukule G, van Asten L, Pereira da Silva S, Aungkulanon S, Bu BJGSI associated MCNetwork. Estimates of global seasonal influenza-associated respiratory mortality: a modelling study. Lancet. 2018;391(10127):1285–300.

2. Bright RA, Medina MJ, Xu X, Perez-Oronoz G, Wallis TR, Davis XM, et al. Incidence of adamantane resistance among influenza A (H3N2) viruses isolated worldwide from 1994 to 2005: A cause for concern. Lancet. 2005;366(9492):1175–81.

3. Deyde VM, Xu X, Bright RA, Shaw M, Smith CB, Zhang Y, et al. Surveillance of resistance to adamantanes among influenza A(H3N2) and A(H1N1) viruses isolated worldwide. Journal of Infectious Diseases. 2007;196(2):249–57.

4. Hayden FG, Sugaya N, Hirotsu N, Lee N, de Jong MD, Hurt AC, et al. Baloxavir Marboxil for Uncomplicated Influenza in Adults and Adolescents. New England Journal of Medicine. 2018;379(10):913–23.

5. Stephenson I, Democratis J, Lackenby A, McNally T, Smith J, Pareek M, et al. Neuraminidase inhibitor resistance after oseltamivir treatment of acute influenza A and B in children. Clinical Infectious Diseases. 2009;48(4):389–96.

6. Petrie JG, Malosh RE, Cheng CK, Ohmit SE, Martin ET, Johnson E, et al. The Household Influenza Vaccine Effectiveness Study: Lack of Antibody Response and Protection Following Receipt of 2014-2015 Influenza Vaccine. Clinical Infectious Diseases. 2017;65(10):1644–51.

7. Kelvin DJ, Farooqui A. Extremely low vaccine effectiveness against influenza H3N2 in the elderly during the 2012/2013 flu season. J Infect Dev Ctries. 2013;7(3):299–301.

8. Tenforde MW, Kondor RJG, Chung JR, Zimmerman RK, Nowalk MP, Jackson ML, et al. Effect of Antigenic Drift on Influenza Vaccine Effectiveness in the United States-2019-2020. Clin Infect Dis. 2021;73(11):e4244–50.

9. Altman MO, Angel M, Košík I, Trovão NS, Zost SJ, Gibbs JS, et al. Human influenza a virus hemagglutinin glycan evolution follows a temporal pattern to a glycan limit. mBio. 2019;10(2):1–15.

10. Kirkpatrick E, Qiu X, Wilson PC, Bahl J, Krammer F. The influenza virus hemagglutinin head evolves faster than the stalk domain. Sci Rep. 2018;8(1):1–14.

11. Heaton NS, Sachs D, Chen CJ, Hai R, Palese P. Genome-wide mutagenesis of influenza virus reveals unique plasticity of the hemagglutinin and NS1 proteins. Proc Natl Acad Sci U S A. 2013;110(50):20248–53.

12. Doud MB, Bloom JD. Accurate measurement of the effects of all amino-acid mutations on influenza hemagglutinin. Viruses. 2016;8(6):1–17.

13. Wang F, Wan Z, Wang Y, Wu J, Fu H, Gao W, et al. Identification of Hemagglutinin Mutations Caused by Neuraminidase Antibody Pressure. Microbiol Spectr. 2021;9(3).

14. Myers JL, Wetzel KS, Linderman SL, Li Y, Sullivan CB, Hensley SE. Compensatory Hemagglutinin Mutations Alter Antigenic Properties of Influenza Viruses. J Virol. 2013;87(20):11168–72.

15. Nachbagauer R, Choi A, Izikson R, Cox MM, Palese P, Krammer F. Age dependence and isotype specificity of influenza virus hemagglutinin stalk-reactive antibodies in humans. mBio. 2016;7(1):1–10.

16. Margine I, Hai R, Albrecht RA, Obermoser G, Harrod AC, Banchereau J, et al. H3N2 Influenza Virus Infection Induces Broadly Reactive Hemagglutinin Stalk Antibodies in Humans and Mice. J Virol. 2013;87(8):4728–37.

17. Moody MA, Zhang R, Walter EB, Woods CW, Ginsburg GS, McClain MT, et al. H3N2 influenza infection elicits more cross-reactive and less clonally expanded anti-hemagglutinin antibodies than influenza vaccination. PLoS One. 2011;6(10).

18. Markham A. Dostarlimab: First Approval. Drugs. 2021;81(10):1213–9.

19. Nowak AK, Cook AM, McDonnell AM, Millward MJ, Creaney J, Francis RJ, et al. A phase 1b clinical trial of the CD40-activating antibody CP-870,893 in combination with cisplatin and pemetrexed in malignant pleural mesothelioma. Annals of Oncology. 2015;26(12):2483–90.

20. Olchanski N, Hansen RN, Pope E, D’Cruz B, Fergie J, Goldstein M, et al. Palivizumab prophylaxis for respiratory syncytial virus: Examining the evidence around value. Open Forum Infect Dis. 2018;5(3):1–9.

21. Weinreich DM, Sivapalasingam S, Norton T, Ali S, Gao H, Bhore R, et al. REGN-COV2, a Neutralizing Antibody Cocktail, in Outpatients with Covid-19. New England Journal of Medicine. 2021;384(3):238–51.

22. Wei G, Meng W, Guo H, Pan W, Liu J, Peng T, et al. Potent neutralization of influenza a virus by a single-domain antibody blocking M2 ion channel protein. PLoS One. 2011;6(12).

23. Cardoso FM, Ibanez LI, Van den Hoecke S, De Baets S, Smet A, Roose K, et al. Single-Domain Antibodies Targeting Neuraminidase Protect against an H5N1 Influenza Virus Challenge. J Virol. 2014;88(15):8278–96.

24. Tillib S V., Ivanova TI, Vasilev LA, Rutovskaya M V., Saakyan SA, Gribova IY, et al. Formatted single-domain antibodies can protect mice against infection with influenza virus (H5N2). Antiviral Res. 2013;97(3):245–54.

25. De Vlieger D, Hoffmann K, Van Molle I, Nerinckx W, Van Hoecke L, Ballegeer M, et al. Selective Engagement of FcγRIV by a M2e-Specific Single Domain Antibody Construct Protects Against Influenza A Virus Infection. Front Immunol. 2019;10.

26. Van Hoecke L, Verbeke R, De Vlieger D, Dewitte H, Roose K, Van Nevel S, et al. mRNA Encoding a Bispecific Single Domain Antibody Construct Protects against Influenza A Virus Infection in Mice. Mol Ther Nucleic Acids. 2020;20(June):777–87.

27. Laursen NS, Friesen RHE, Zhu X, Jongeneelen M, Blokland S, Vermond J, et al. Universal protection against influenza infection by a multidomain antibody to influenza hemagglutinin. Science (1979). 2018;362(6414):598–602.

28. Hufton SE, Risley P, Ball CR, Major D, Engelhardt OG, Poole S. The breadth of cross sub-type neutralisation activity of a single domain antibody to influenza hemagglutinin can be increased by antibody valency. PLoS One. 2014;9(8).

29. Del Rosario JMM, Smith M, Zaki K, Risley P, Temperton N, Engelhardt OG, et al. Protection From Influenza by Intramuscular Gene Vector Delivery of a Broadly Neutralizing Nanobody Does Not Depend on Antibody Dependent Cellular Cytotoxicity. Front Immunol. 2020;11(May):1–15.

30. Gaiotto T, Hufton SE. Cross-neutralising nanobodies bind to a conserved pocket in the hemagglutinin stem region identified using yeast display and deep mutational scanning. PLoS One. 2016;11(10):1–27.

31. Conrath KE, Lauwereys M, Galleni M, Matagne A, Frère JM, Kinne J, et al. β-Lactamase inhibitors derived from single-domain antibody fragments elicited in the Camelidae. Antimicrob Agents Chemother. 2001;45(10):2807–12.

32. Lauwereys M, Ghahroudi MA, Desmyter A, Kinne J, Hölzer W, De Genst E, et al. Potent enzyme inhibitors derived from dromedary heavy-chain antibodies. EMBO Journal. 1998;17(13):3512–20.

33. Jank L, Pinto-Espinoza C, Duan Y, Koch-Nolte F, Magnus T, Rissiek B. Current Approaches and Future Perspectives for Nanobodies in Stroke Diagnostic and Therapy. Antibodies. 2019;8(1):5.

34. Jegaskanda S, Weinfurter JT, Friedrich TC, Kent SJ. Antibody-Dependent Cellular Cytotoxicity Is Associated with Control of Pandemic H1N1 Influenza Virus Infection of Macaques. J Virol. 2013;87(10):5512–22.

35. Asthagiri Arunkumar G, Ioannou A, Wohlbold TJ, Meade P, Aslam S, Amanat F, et al. Broadly Cross-Reactive, Nonneutralizing Antibodies against Influenza B Virus Hemagglutinin Demonstrate Effector Function-Dependent Protection against Lethal Viral Challenge in Mice. J Virol. 2019;93(6):1–17.

36. Henry Dunand CJ, Leon PE, Huang M, Choi A, Chromikova V, Ho IY, Tan GS, Cruz J, Hirsh A, Zheng NY, Mullarkey CE, Ennis FA, Terajima M, Treanor JJ, Topham DJ, Subbarao K, Palese P, Krammer F WPC. Both Neutralizing and Non-neutralizing Human H7N9 Influenza Vaccine-induced Monoclonal Antibodies Confer Protection. Cell Host Microbe. 2016;19(6):800–13.

37. DiLillo DJ, Tan GS, Palese P RJV. Broadly neutralizing hemagglutinin stalk–specific antibodies require FcγR interactions for protection against influenza virus in vivo. Nat Med. 2014;20(2):143–51.

38. De Vries RD, Nieuwkoop NJ, Van Der Klis FRM, Koopmans MPG, Krammer F, Rimmelzwaan GF. Primary human influenza B virus infection induces cross-lineage hemagglutinin stalk-specifc antibodies mediating antibody-dependent cellular cytoxicity. Journal of Infectious Diseases. 2018;217(1):3–11.

39. He W, Tan GS, Mullarkey CE, Lee AJ, Lam MMW, Krammer F, et al. Epitope specificity plays a critical role in regulating antibody-dependent cell-mediated cytotoxicity against influenza A virus. Proc Natl Acad Sci U S A. 2016;113(42):11931–6.

40. Vanderven HA, Kent SJ. The protective potential of Fc-mediated antibody functions against influenza virus and other viral pathogens. Immunol Cell Biol. 2020;98(4):253–63.

41. Short KR, Kroeze EJBV, Fouchier RAM, Kuiken T. Pathogenesis of influenza-induced acute respiratory distress syndrome. Lancet Infect Dis. 2014;14(1):57–69.

42. Respaud R, Vecellio L, Diot P, Heuzé-Vourc’h N. Nebulization as a delivery method for mAbs in respiratory diseases. Expert Opin Drug Deliv. 2015;12(6):1027–39.

43. Leyva-Grado VH, Tan GS, Leon PE, Yondola M, Palese P. Direct administration in the respiratory tract improves efficacy of broadly neutralizing anti-influenza virus monoclonal antibodies. Antimicrob Agents Chemother. 2015;59(7):4162–72.

44. Vigil A, Frias-Staheli N, Carabeo T, Wittekind M. Airway Delivery of Anti-influenza Monoclonal Antibodies Results in Enhanced Antiviral Activities and Enables Broad-Coverage Combination Therapies. J Virol. 2020;94(22):1–20.

45. Wu X, Cheng L, Fu M, Huang B, Zhu L, Xu S, et al. A potent bispecific nanobody protects hACE2 mice against SARS-CoV-2 infection via intranasal administration. Cell Rep. 2021;37(3):109869.

46. Wu X, Wang Y, Cheng L, Ni F, Zhu L, Ma S, et al. Short-Term Instantaneous Prophylaxis and Efficient Treatment Against SARS-CoV-2 in hACE2 Mice Conferred by an Intranasal Nanobody (Nb22). Front Immunol. 2022;13(March):1–15.

47. S Raut, J Hurd, G Blandford, R B Heath RJC. The pathogenesis of infections of the mouse caused by virulent and avirulent variants of an influenza virus. 1975;8(1):127–36.

48. Voronina D V., Shcheblyakov D V., Esmagambetov IB, Derkaev AA, Popova O, Shcherbinin DN. Development of Neutralizing Nanobodies to the Hemagglutinin Stem Domain of Influenza A Viruses. Acta Naturae. 2021;13(4):33–41.

49. Chiang YW, Li CJ, Su HY, Hsieh KT, Weng CW, Chen HW, et al. Development of mouse monoclonal antibody for detecting hemagglutinin of avian influenza A(H7N9) virus and preventing virus infection. Appl Microbiol Biotechnol. 2021;105(8):3235–48.

50. Hong Y, Guo H, Wei M, Zhang Y, Fang M, Cheng T, et al. Cell-based reporter assays for measurements of antibody-mediated cellular cytotoxicity and phagocytosis against SARS-CoV-2 spike protein. J Virol Methods. 2022;307(January):293.

51. Reed, L.J. and Muench H. A Simple Method of Estimating Fifty Percent Endpoints. 1938;27:493–7.

52. Kaufmann B, Nybakken GE, Chipman PR, Zhang W, Diamond MS, Fremont DH, et al. West Nile virus in complex with the Fab fragment of a neutralizing monoclonal antibody. Proc Natl Acad Sci U S A. 2006;103(33):12400–4.

53. Thouvenin E, Hewat E. When two into one won’t go: Fitting in the presence of steric hindrance and partial occupancy. Acta Crystallogr D Biol Crystallogr. 2000;56(10):1350–7.

54. DiLillo DJ, Palese P, Wilson PC, Ravetch J V. Broadly neutralizing anti-influenza antibodies require Fc receptor engagement for in vivo protection. Journal of Clinical Investigation. 2016;126(2):605–10.

55. Sun H, Xiao Y, Liu J, Wang D, Li F, Wang C, et al. Prevalent Eurasian avian-like H1N1 swine influenza virus with 2009 pandemic viral genes facilitating human infection. Proc Natl Acad Sci U S A. 2020;117(29):17204–10.

56. Pappas C, Yang H, Carney PJ, Pearce MB, Katz JM, Stevens J TT. Assessment of transmission, pathogenesis and adaptation of H2 subtype influenza viruses in ferrets. Virology. 2015;477:61–71.

57. Sutton TC. The pandemic threat of emerging H5 and H7 avian influenza viruses. Viruses. 2018;10(9):1–21.

58. Song W, Qin K. Human-infecting influenza A (H9N2) virus: A forgotten potential pandemic strain? Zoonoses Public Health. 2020;67(3):203–12.

59. Vanderven HA, Liu L, Ana-Sosa-Batiz F, Nguyen THO, Wan Y, Wines B, et al. Fc functional antibodies in humans with severe H7N9 and seasonal influenza. Journal of Clinical Investigation. 2017;2(13):1–15.

60. Rotman M, Welling MM, van den Boogaard ML, Moursel LG, van der Graaf LM, van Buchem MA, et al. Fusion of hIgG1-Fc to 111In-anti-amyloid single domain antibody fragment VHH-pa2H prolongs blood residential time in APP/PS1 mice but does not increase brain uptake. Nucl Med Biol. 2015;42(8):695–702.

61. Wu B, Sun YN. Pharmacokinetics of peptide-Fc fusion proteins. J Pharm Sci. 2014;103(1):53–64.

62. Schepens B, van Schie L, Nerinckx W, Roose K, Van Breedam W, Fijalkowska D, et al. An affinity-enhanced, broadly neutralizing heavy chain-only antibody protects against SARS-CoV-2 infection in animal models. Sci Transl Med. 2021;13(621):1–18.

63. Lee PS, Yoshida R, Ekiert DC, Sakai N, Suzuki Y, Takada A, et al. Heterosubtypic antibody recognition of the influenza virus hemagglutinin receptor binding site enhanced by avidity. Proc Natl Acad Sci U S A. 2012;109(42):17040–5.

64. Ekiert DC, Bhabha G, Elsliger M andré, Friesen RHE, Jongeneelen M, Throsby M, et al. Antibody recognition of a highly conserved influenza virus epitope: implications for universal prevention and therapy. Science (1979). 2009;324(5924):246–51.

65. Throsby M, van den Brink E, Jongeneelen M, Poon LLM, Alard P, Cornelissen L, et al. Heterosubtypic neutralizing monoclonal antibodies cross-protective against H5N1 and H1N1 recovered from human IgM+ memory B cells. PLoS One. 2008;3(12).

66. Dreyfus C, Laursen NS, Kwaks T, Zuijdgeest D, Khayat R, Ekiert DC, et al. Highly conserved protective epitopes on influenza B viruses. Science (1979). 2012;337(6100):1343–8.

67. Parray HA, Shukla S, Perween R, Khatri R, Shrivastava T, Singh V, et al. Inhalation monoclonal antibody therapy: a new way to treat and manage respiratory infections. Appl Microbiol Biotechnol. 2021;105(16–17):6315–32.

68. He F, Kumar SR, Syed Khader SM, Tan Y, Prabakaran M, Kwang J. Effective intranasal therapeutics and prophylactics with monoclonal antibody against lethal infection of H7N7 influenza virus. Antiviral Res. 2013;100(1):207–14.

69. Prabakaran M, Prabhu N, He F, Hongliang Q, Ho HT, Qiang J, et al. Combination therapy using chimeric monoclonal antibodies protects mice from lethal H5N1 infection and prevents formation of escape mutants. PLoS One. 2009;4(5).

70. Baum A, Ajithdoss D, Copin R, Zhou A, Lanza K, Negron N, et al. REGN-COV2 antibodies prevent and treat SARS-CoV-2 infection in rhesus macaques and hamsters. Science (1979). 2020;370(6520):1110–5.

71. Pascal KE, Dudgeon D, Trefry JC, Anantpadma M, Sakurai Y, Murin CD, et al. Development of Clinical-Stage Human Monoclonal Antibodies That Treat Advanced Ebola Virus Disease in Nonhuman Primates. Journal of Infectious Diseases. 2018;218(Suppl 5):S612–26.

72. Yi KS, Choi JA, Kim P, Ryu DK, Yang E, Son D, et al. Broader neutralization of CT-P27 against influenza A subtypes by combining two human monoclonal antibodies. PLoS One. 2020;15(7 July):1–14.

73. Sutton TC, Lamirande EW, Bock KW, Moore IN, Koudstaal W, Rehman M, et al. In Vitro Neutralization Is Not Predictive of Prophylactic Efficacy of Broadly Neutralizing Monoclonal Antibodies CR6261 and CR9114 against Lethal H2 Influenza Virus Challenge in Mice. J Virol. 2017;91(24):1–12.

74. Yamayoshi S, Uraki R, Ito M, Kiso M, Nakatsu S, Yasuhara A, et al. A Broadly Reactive Human Anti-hemagglutinin Stem Monoclonal Antibody That Inhibits Influenza A Virus Particle Release. EBioMedicine. 2017;17:182–91.

75. Kajihara M, Marzi A, Nakayama E, Noda T, Kuroda M, Manzoor R, et al. Inhibition of Marburg Virus Budding by Nonneutralizing Antibodies to the Envelope Glycoprotein. J Virol. 2012;86(24):13467–74.

76. Dufloo J, Planchais C, Frémont S, Lorin V, Guivel-Benhassine F, Stefic K, et al. Broadly neutralizing anti-HIV-1 antibodies tether viral particles at the surface of infected cells. Nat Commun. 2022;13(1):1–11.

77. Kosik I, Angeletti D, Gibbs JS, Angel M, Takeda K, Kosikova M, et al. Neuraminidase inhibition contributes to influenza A virus neutralization by anti-hemagglutinin stem antibodies. Journal of Experimental Medicine. 2019;216(2):304–16.

78. Wohlbold TJ, Chromikova V, Tan GS, Meade P, Amanat F, Comella P, et al. Hemagglutinin Stalk- and Neuraminidase-Specific Monoclonal Antibodies Protect against Lethal H10N8 Influenza Virus Infection in Mice. J Virol. 2016;90(2):851–61.

